# Testing Equality of Curves after Covariate Adjustment

**DOI:** 10.1101/228148

**Authors:** Parichoy Pal Choudhury, Prosenjit Kundu, Qian-Li Xue, Reyhan Westbrook, Ciprian M. Crainiceanu

## Abstract

This paper is concerned with providing simple methodological approaches for global and local tests of difference between the mean of treatment and control groups when the measured outcome is a function. The added complexity is that for every subject we have repeated samples for the same curve and additional covariates of interest. We propose a permutation based approach to test for a global difference between the averages of two functional processes after covariate adjustment. The within group averages are estimated by modeling the relationship of the functional outcome on the covariate using functional regression methods and then averaging with respect to the covariate distribution in each group. The test statistic is the *L*^2^ area under the squared difference curve. We also test for the localized differences between the two average curves using a nonparametric bootstrap of subjects to obtain the 95% pointwise and joint confidence intervals for the estimated covariate-adjusted difference curve. Extensive simulation studies illustrate that the proposed tests preserve the type one error and are highly sensitive to detecting departures from the null assumption. We illustrate our method by studying the differences in time varying oxygen consumption between the frail Interleukin 10tm1Cgn (IL10tm) mice and the wildtype mice after adjusting for body composition measures.

## 1. Introduction

We propose simple methodological approaches for global and local tests of difference between the mean of treatment and control groups when the measured outcome is a function. Several papers in the functional data analysis literature have focussed on comparing the averages of two functional processes. For example, Benko et al. (2009) developed bootstrap-based tests of equality of means, eigenvalues and eigenfunctions of covariance function in the two sample problem. Hall and Van Keilegom (2007) used bootstrap-based tests for equality of distributions of two independent samples of curves. Zhang et al. (2010) proposed *L*^2^- based and bootstrap-based statistics for testing equality of two average curves when the subject specific curves are independent and observed without noise. Crainiceanu et al. (2012) proposed a bootstrap-based inference procedure for the difference in means of two correlated functional processes. However, neither of these approaches considered covariate-adjusted testing, which is essential in cases when covariates may differ across groups. Several authors have developed Bayesian approaches for this problem that allow for covariate adjustment and model the individual curves as well as possible correlations between them. (Behseta and Kass, 2005; Behseta et al., 2007; Morris et al., 2003; Morris and Carroll, 2006; Morris et al., 2008, 2011). In this work, we focus on developing simple test procedures based on the estimand of interest directly.

The scientific problem that motivated our study is whether the targeted deletion of the interleukin 10 gene (IL-10^*tm*1*Cgn*^) in mice leads to decrease in oxygen consumption. We have repeated measures of oxygen consumption in a group of approximately 22 month-old mice. In total, there were 20 animals and the gene was knocked out in 10 of them (IL 10 -/- group) and present in the others (control group). Oxygen consumption measurements were taken for each mouse at regularly spaced time points over four days. We want to explore if the average oxygen consumption through the day (midnight-midnight) differs significantly between the two groups and if the genotype-outcome association is altered by the body composition of the animal. The novelty of our approach is that it addresses the problem that each animal has repeated functional measurements over multiple days (oxygen consumption measured every 30 minutes for 4 days) and additional covariates of interest (i.e., body composition measures).

We develop a permutation based approach to test for a global difference between the averages of two functional processes after covariate adjustment using the estimated *L*^2^ area under the squared difference curve as the test statistic. We also test for localized differences between the two covariate adjusted average curves using the 95% pointwise and joint confidence intervals obtained using a nonparametric bootstrap of subjects. The main novelty of our paper is that we are using the covariate adjusted curves to develop the test procedures and take into account the within-subject sampling functional correlation. The proposed approach is easy-to-implement, computationally fast, scalable and adaptable to more complex settings. In Section 2 we develop the statistical framework for our method. Section 3 provides the results of the real data analysis and the simulation study. We conclude with a discussion in Section 4.

## 2. Methods

Our method utilizes information from “treated” animals (e.g., IL10^*tm*^ group) and animals who are “not treated” (e.g., control group). Both groups have information on a functional outcome [e.g., oxygen consumption measured at regular intervals (30 minutes) over a period of time (4 days)] and baseline covariates (e.g., body composition measures). Figure 1 displays the scatter plots for oxygen consumption of the animals in each group in a 24 hour period (midnight-midnight) over 4 days. Figure 2 displays the average oxygen consumption (over every observation within days and all four days) of each subject as a function of body mass composition as well as the body mass composition distribution within treatment groups. Let *n_i_* denote the number of animals in the *i^th^* group, *i* = 0 for “treated” animals and *i* = 1 for “not treated” animals. We observe {(*Y_ijl_*(*t*),*X_ij_*): *t* = *t*_1_,…,*t_k_*; *i* = 0,1; *j* = 1,…, *n_i_*; l = 1, 2, 3, 4}, where *Y_ijl_*(*t*) denotes the functional outcome observed at the time points *t*_1_,…, *t_k_* in the range [0,*T*] during the *l^th^* day and *X_ij_* denotes the vector of baseline covariates for the *j^th^* animal in the *i^th^* group. Denote by 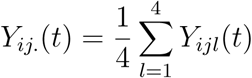. We are interested in a model of the type:

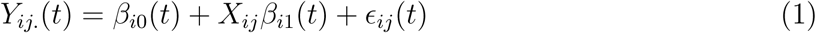

where є_ij_(*t*) is a mean zero process with unspecified correlation structure. We want to test the hypothesis: *μ*_1_(*t*) = *μ*_0_(*t*), where *μ_i_*(*t*) = *E*[*Y*_*ij*._(*t*)] for *i* = 0,1. Note that *μ_i_*(*t*) = *E*[*Y_ij._*(*t*)] = *E*[*E*[*Y_ij._*(*t*)|*X_ij_*]] = *β*_i0_(*t*) + *β_i_*_1_(*t*)*E*[*X_ij_*]. We model the functional regression parameters in equation 1 using P-splines that combine a B-spline basis with a discrete penalty on the basis coefficients (Eilers et al., 1996). The regression functions are estimated using restricted maximum likelihood estimation of the associated penalized least squares objective function in the framework of generalized additive models (Chambers et al., 1991; Hastie and Tibshirani, 1990). For *i* = 0,1 we estimate *μ_i_*(*t*) by 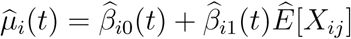, where 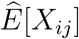 is the sample average of the covariates in the *i^th^* group. We are making the working assumption that є_*ij*_(*t*) are independent. This assumption substantially simplifies the estimation procedure, though for inference, the within-subject correlation will need to be taken into account. Estimating parameters under independence and then correcting the confidence intervals has a long and successful history in Statistics.

**Figure 1.**
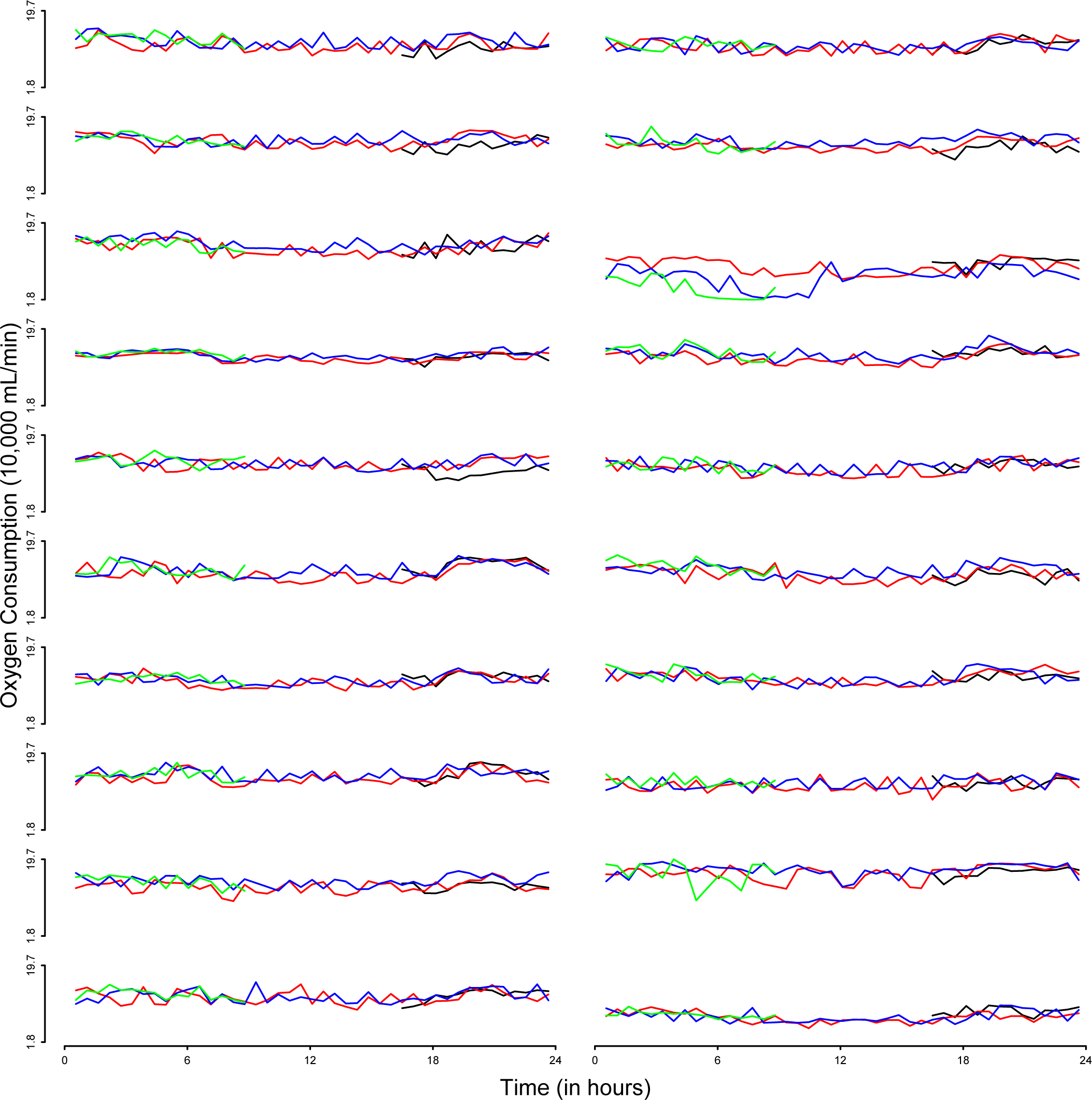
Plot of oxygen consumption during the 24 hours over multiple days; the panels in the left correspond to animals in the control group and panels on the right correspond to animals in the IL-10^*tm*1*Cgn*^ group. Each panel shows the oxygen consumption of an animal over 4 days: Day 1 (black line), Day 2 (red line), Day 3 (blue line) and Day 4 (green line).

**Figure 2.**
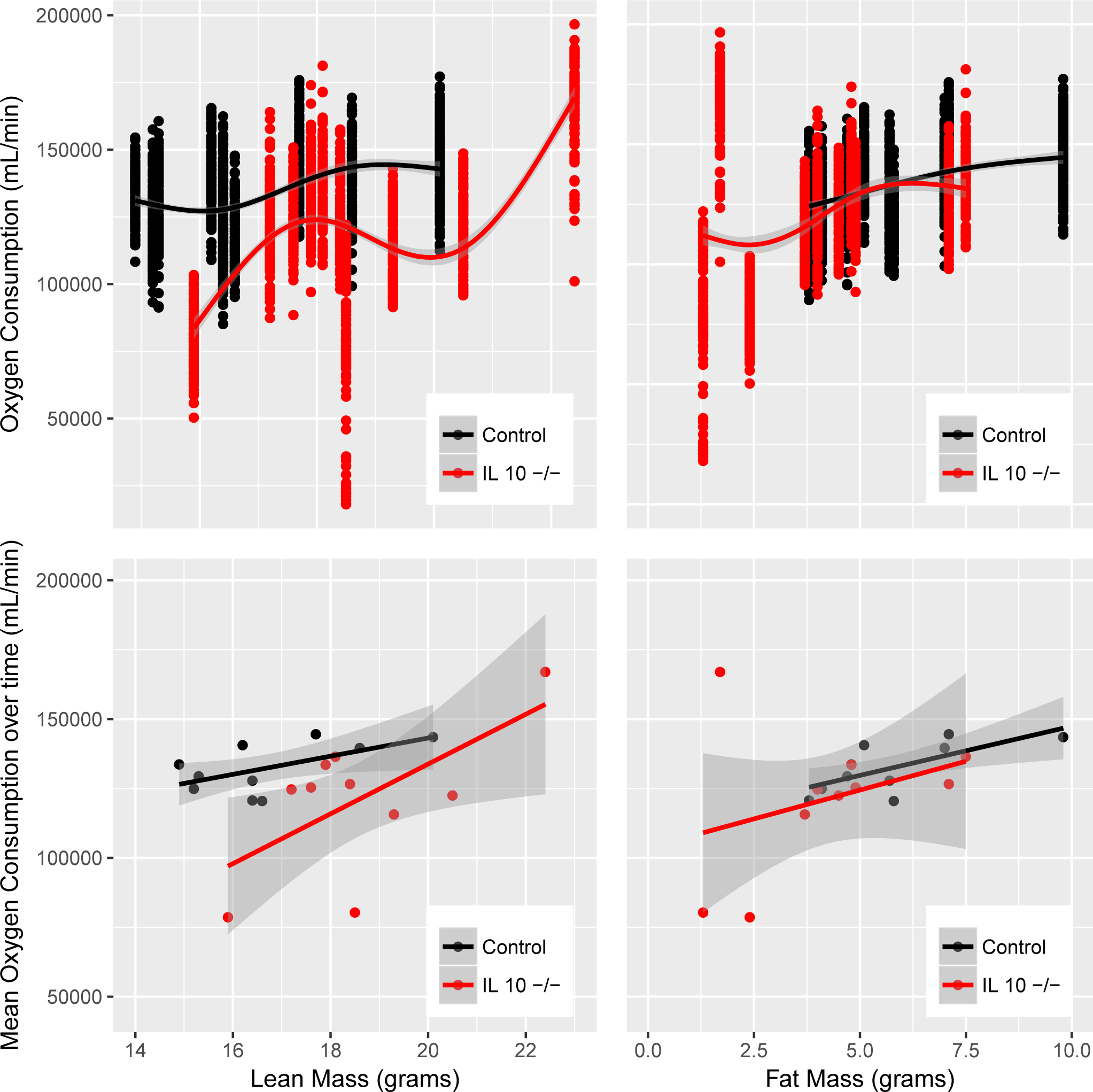
Bivariate relationship of oxygen consumption with lean mass and fat mass in the two genotype groups: in the upper panel the dependent variable is observed oxygen consumption at all time points; in the lower panel the dependent variable is average oxygen consumption over time

### 2.1 Test for Global Difference

Define *δ*(*t*) = *μ*_1_(*t*) – *μ*_0_(*t*) and denote by 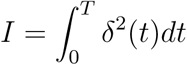 Note that *δ*(*t*) = 0 for every *t* if and only if *I* = 0. Thus *I* is a measure of global difference between the means of the two groups.

Denote the estimates of the within group averages by 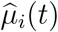, *i* = 0,1 and let 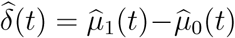 be an estimator of *δ*(*t*). We estimate *I* by 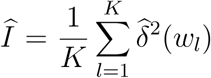 where w_1_,…, w_*K*_ is a fine grid [0,*T*]. We consider the null hypothesis: *H*_0_: *I* = 0 versus the alternative hypothesis *H_a_*: *I* > 0. Our permutation based test procedure involves the following steps:

i. Consider the joint dataset with *n* = *n*_1_ + *n*_0_ animals, where for *i* = 0,1, *n_i_* animals come from *i^th^* group. Consider a random permutation *p* of the labels of “treatment” (i.e. “treated” or “not treated”).
ii. For each permuted dataset, p, estimate the averages of the two groups: 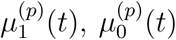 by the model fitting and estimation procedure described earlier. Denote the difference function 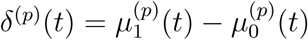 and the integral of the squared difference function by *I_p_*. Compute 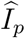 using the method described earlier.
iii. Repeat step (i) with *P* permuted datasets.
iv. Compute the permutation test p-value to be the proportion of permutations with 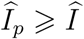. The key idea of the permutation test is as follows: under the null hypothesis there is no difference in the average outcome between the two groups. Hence the treatment labels are exchangeable under the null hypothesis. The empirical distribution of 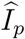 estimates the distribution of the global difference between the two groups under the null hypothesis. This provides the rationale for the computation of the test p-value in step (iv).

### 2.2 Test for Localized Differences

We also propose a test for localized differences between groups, (i.e. the difference in average outcome at particular time points) using a nonparametric bootstrap-based inferential procedure (Crainiceanu et al., 2012). The main difference from the procedure in Crainiceanu et al. (2012) is that we are working with covariate adjusted curves, which is important in many applications.

One question of interest is whether there is a difference in the average outcomes between the groups at a fixed time point *t*. The corresponding null and alternative hypotheses can be stated as:

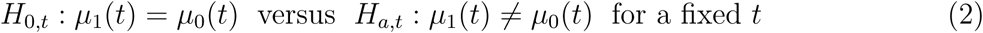

We compute the 95% pointwise confidence intervals to address this question.

Another question of interest is whether there is a difference between the average curves at all time points. The corresponding null and alternative hypotheses are as follows:

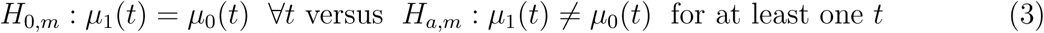

This question can be addressed using the joint confidence intervals to account for multiple hypotheses testing.

The key steps in the computation of the different kind of confidence intervals are as follows:

i. Generate *B* simple random samples with replacement separately from each group.
ii. For each bootstrap dataset, define *δ_b_*(*t*) = *μ*_1*b*_(*t*) – *μ*_0*b*_(*t*), *b* = 1,…,*B*. Estimate *μ*_1*b*_(*t*), *μ*_0*b*_(*t*) and *δ_b_*(*t*) using the procedure described earlier.
iii. Compute 95% pointwise and joint confidence intervals.

The 95% pointwise confidence intervals in step (iii) are constructed based on the bootstrap distribution of *δ_b_*(*t*) for a fixed *t*. They can be interpreted as follows: at each time point *t* in bootstrapped samples the true difference will be covered by the interval 95% of the time. The 95% joint confidence intervals are computed by the algorithm given in Section 3 of Crainiceanu et al. (2012). The interpretation is as follows: at all time points in repeated samples the true difference will be covered by the interval 95% of the time.

## 3. Results

A mouse with targeted deletion in the interleukin 10 gene (IL-10^*tm*1*Cgn*^) has been proposed as a mouse model for frailty and low-grade inflammation. The older frail IL-10*^*tm*^* mice show many similarities with older frail human beings (Sikka, 2013). This provides the rationale for using it as a scientific model for studying frailty. We use the methods developed in the Section 2 to explore if older frail IL-10^*tm*^ mice have reduced oxygen consumption compared to normal wildtype mice. We further investigate whether the statistical association between the genotype and decreased oxygen consumption is altered by the body composition of the animal.

### 3.1 Description of study design and data

We have experimental data on *n*_0_ = 10 mice with the interleukin 10 gene knocked out (IL-10^*tm*^ group) and *n*_1_ = 10 additional mice where the gene is present (control group). For the animals in each group we have repeated measures of oxygen consumption per gram body weight (every 30 mins over 4 consecutive days; 116 repeated observations for each animal, cf Figure 1). We also have information on body composition measures (body weight, lean mass, fat mass, fluid mass) for each animal obtained through Nuclear Magnetic Resonance (NMR) experiments. We want to compare the average daily oxygen consumption curves between the groups after adjusting for body composition measures.

### 3.2 Exploratory Analysis, Outlier Identification and Covariate Adjustment

Figure 2 displays the bivariate distributions of oxygen consumption and the lean mass and fat mass in the two genotype groups. The upper panel includes observations at all time points as the dependent variable. The lower panel uses the average oxygen consumption over time as the dependent variable. For illustrative purposes we used thin plate regression spline smoothing with four basis functions. The two genotype groups show similar relationship between oxygen consumption and fat mass. However, there is a difference in the relationship between oxygen consumption and lean mass across the groups.

Figure 3 displays the relationship between lean mass and fat mass for both groups of animals using a thin plate regression spline smoother with four basis functions. Based on visual inspection, we identified one outlying animal in the IL-10^*tm*^ group with lean mass 22.4 grams and fat mass 1.7 grams. The striking difference in the nature of the red curve in the upper and lower panel of Figure 3 supports this fact. In addition to the main analysis (Analysis I) that includes all the animals, we also perform a sensitivity analysis excluding this outlying animal (Analysis II).

**Figure 3.**
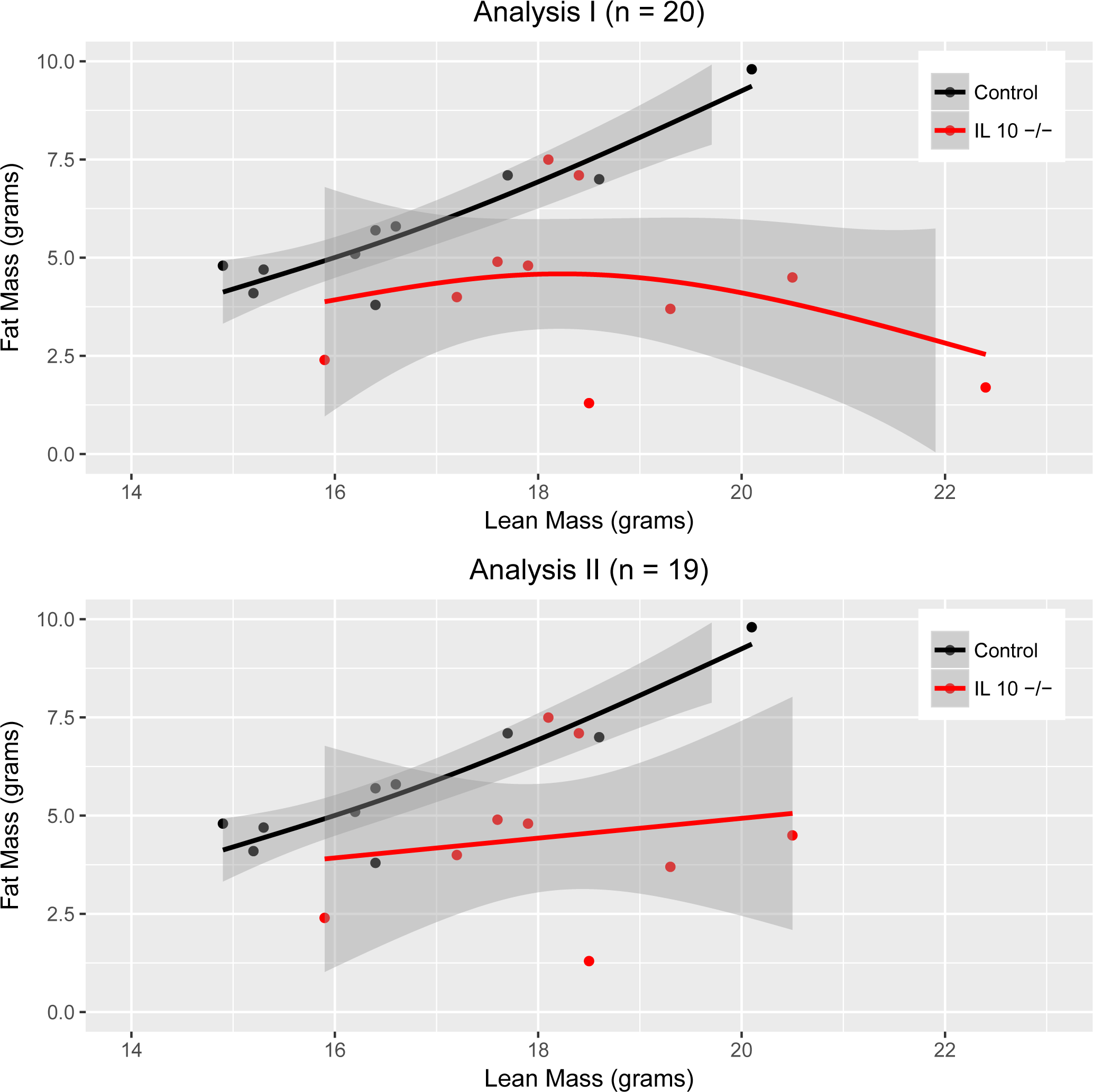
Relationship between lean mass and fat mass in the two genotype groups: upper panel corresponds to Analysis I (*n* = 20, *n*_0_ = 10, *n*_1_ = 10) with the outlier, lower panel corresponds to Analysis II (*n* = 19, *n*_0_ = 9, *n*_1_ = 10) without the outlier.

Figure 4 displays the bivariate distribution of oxygen consumption and the ratio of fat mass and lean mass in the two genotype groups for both Analyses I and II. The upper panel includes observations at all time points as the dependent variable. The lower panel uses the average oxygen consumption over time as the dependent variable. We use thin plate regression splines with four basis functions to smooth the data. The animals in the IL-10^*tm*^ group have lower values of the ratio of fat mass and lean mass compared to the animals in the control group. These plots also indicate that the animal mentioned earlier may be an outlier.

**Figure 4.**
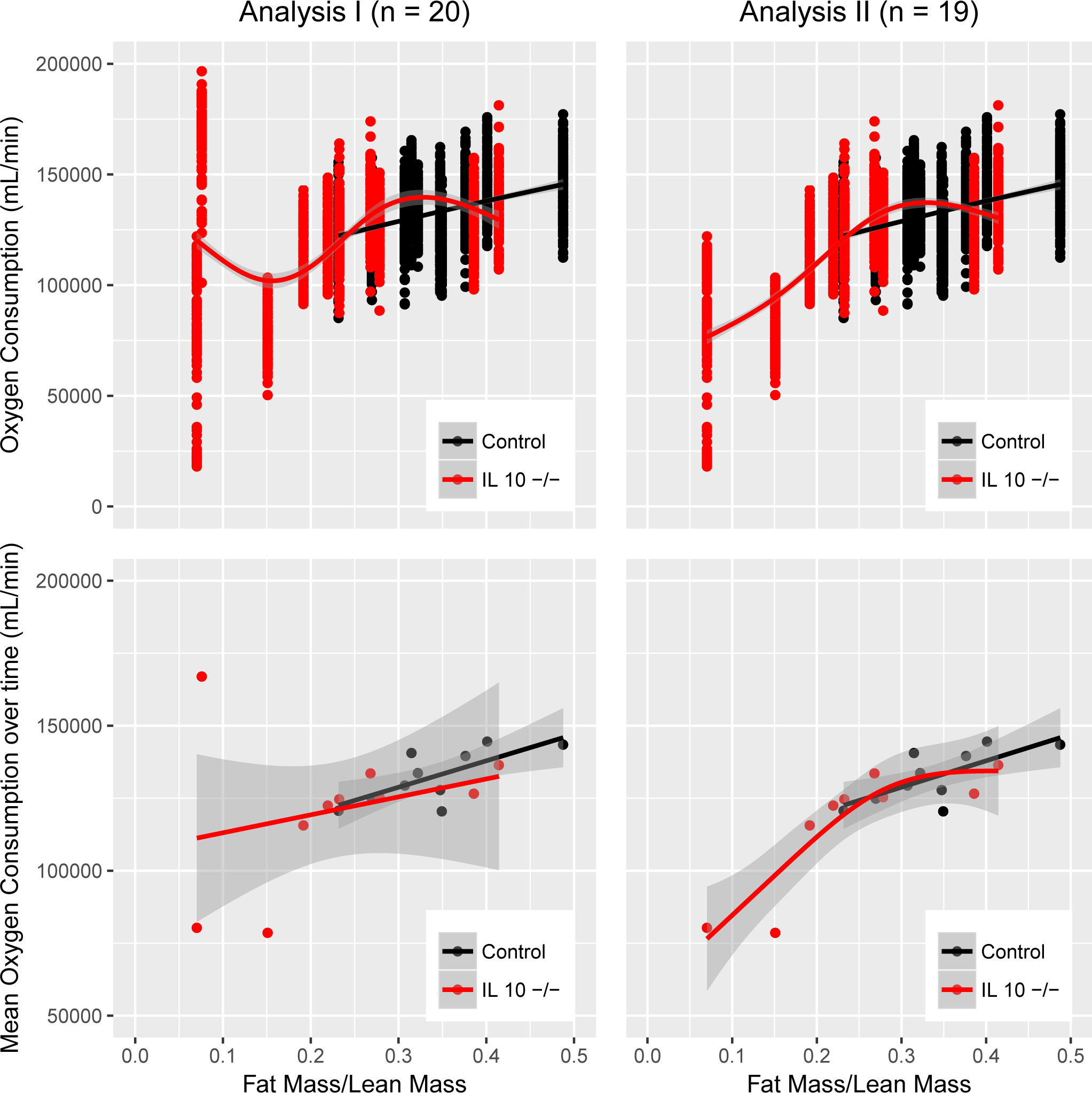
Bivariate relationship of oxygen consumption with ratio of fat mass and lean mass in the two genotype groups for both Analysis I (*n* = 20, *n*_0_ = 10, *n*_1_ = 10) and Analysis II (*n* = 19, *n*_0_ = 9,*n*_1_ = 10): in the upper panel the dependent variable is observed oxygen consumption at all time points; in the lower panel the dependent variable is average oxygen consumption over time.

The exploratory analyses indicate that the lean mass and the ratio of fat mass and lean mass are key body composition measures that could potentially modify the association between genotype and oxygen consumption. For the rest of the analysis, we use the subject-level time-specific oxygen consumption averaged over the four days as our functional outcome and consider lean mass and fat mass to lean mass ratio as covariates for adjustment. The covariate adjusted curves are computed by the methods described in Section 2. We also compare results with the standard approach in the literature that normalizes the outcome by per gram body weight. (Speakman, 2013). To implement this approach, we fit intercept only functional regression models in both the genotype groups and estimate the average oxygen consumption.

### 3.3 Global Genotype Effect

We follow the permutation based approach outlined in Section 2.1 to test for global genotype effect. When we use the body weight normalized outcome the permutation test results show significant global genotype effect (*p-value* = 0.007). However, after adjustment for lean mass and fat mass to lean mass ratio, the effect is no longer significant (*p-value* = 0.19).

### 3.4 Localized Genotype Effect

Figure 5 shows the point estimates and the 95% pointwise and joint confidence intervals for the difference in average oxygen consumption between the control mice and the IL-10^*tm*^ mice. When the oxygen consumption is normalized per gram of body weight (left panel) we observe a significant decrease in average oxygen consumption at different times during the day for the IL-10^*tm*^ mice compared to the control mice. However, the difference is no longer statistically significant after adjustment for lean mass and the ratio of fat mass and lean mass.

**Figure 5.**
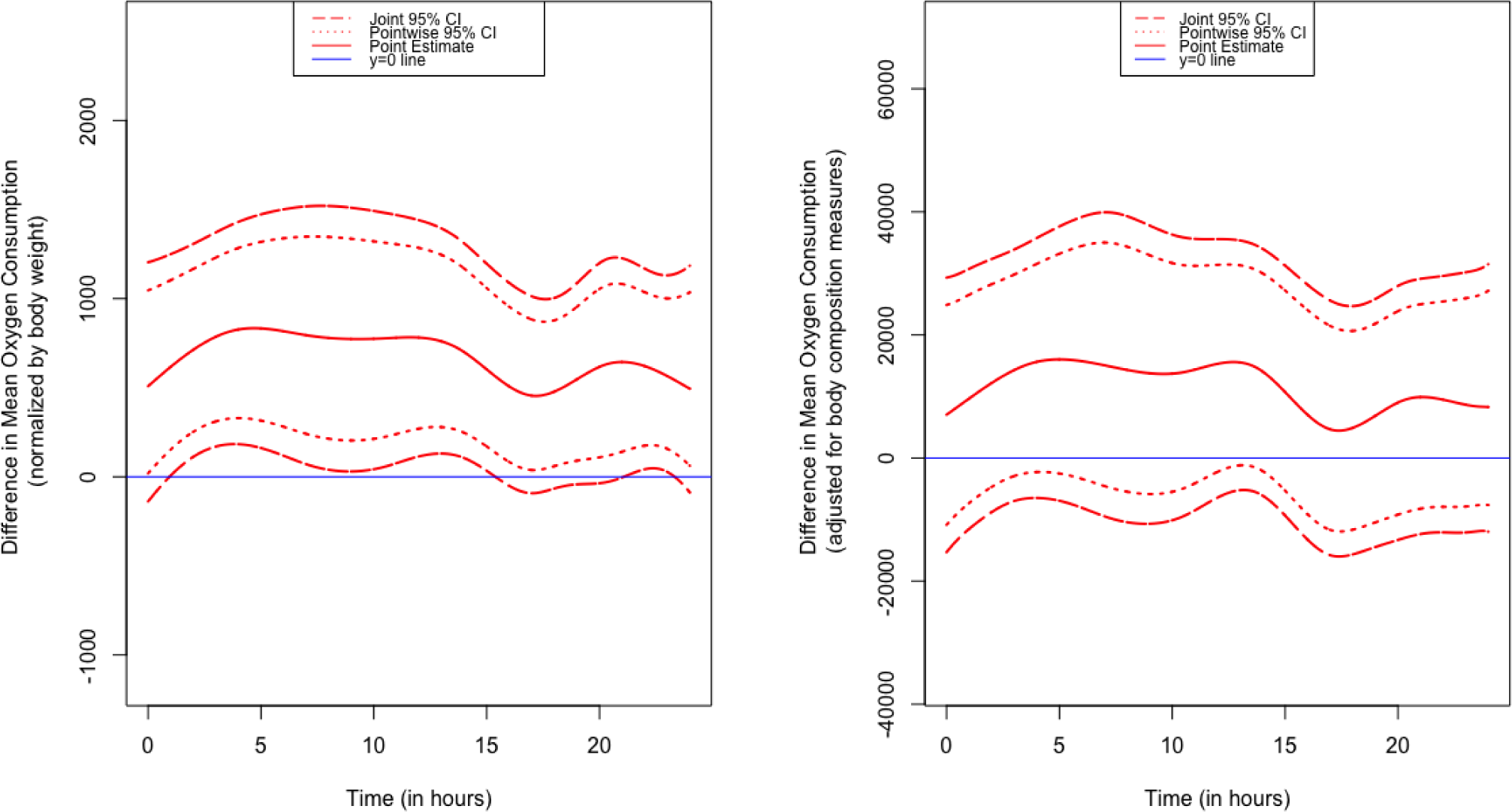
Plots showing 95% pointwise and joint confidence intervals for the difference in mean oxygen consumption between control mice and IL-10^*tm*^ mice: in the left panel the oxygen consumption is normalized by body weight (BW) and in the right panel the oxygen consumption is adjusted for the ratio of fat mass and lean mass (FM/LM) of the animal.

The results show some evidence of genotype effect being mediated through body composition measures. We report the results of the sensitivity analysis (Analysis II) in the the Appendix. The results are sensitive to the outlying animal. The total computation time for performing the tests for global genotype effect and localized genotype effect was around 10 mins (2.8 GHz intel core i7, 16 GB RAM Macbook Pro)

### 3.5 Simulation Results

We investigate the performance of the proposed methods in a simulation study. For different settings we generate 500 datasets from Model 1 with two covariates, for different choices of *β*_*i*0_(*t*) and *β*_*i*1_(*t*). We consider a time grid of 100 equally spaced points in the interval [0,1]. We generate *є*_*ij*_(*t*) in Model 1 from a Gaussian Process distribution characterized by the equation 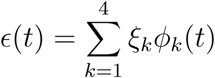 where *ξ_k_* are mutually independent *N*(0, λ*_k_*) for *k* = 1, 2, 3, 4 and λ*_k_* and *ϕ*_*k*_(*t*) represent the *k^th^* eigenvalue and eigenfunction respectively of the functional principal component decomposition of a centered and scaled version of outcome data. We consider different settings that combine the choices of the following parameters:

1. Number of subjects: consider *n*_1_ = *n*_0_ with *n* = n_1_ + *n*_0_ and take *n* = 10, 20, 50,100.
2. We generate the two covariates independently using the same distribution in each group. More specifically, we use the empirical distribution of lean mass and fat mass to lean mass ratio in the control group of the data to generate covariates. For the regression functions consider two scenarios for data generated under the null hypothesis i.e., *δ*(*t*) = 0: (i) *β*_00_(*t*) = *β*_10_(*t*) = 0.91 and *β*_01_(*t*) = *β*_11_(*t*) = (–0.77, 0.04); (ii) *β*_00_(*t*) = *β*_10_(*t*) = sin *πt*, *β*_01_(*t*) = *β*_11_(*t*) = (–0.77, 0.04); two additional scenarios for data generated under the alternative hypothesis i.e., *δ*(*t*) ≠ 0: (iii) *β*_00_(*t*) = 0.91,*β*_10_(*t*) = 1.40, *β*_01_(*t*) = *β*_11_(*t*) = (–0.77, 0.04); (iv) *β*_00_(*t*) = 0.1(1+*t*)^2^,*β*_10_(*t*) = 0.5(1ȁ*t*)^2^, *β*_01_(*t*) = *β*_11_(*t*) = (–0.77, 0.04). The regression function for the covariates were chosen as the average (over time) of the estimated regression functions based on fitting model 1 on the data.

For the test of global differences, let *P* be the conditional probability that the null hypothesis is rejected given the true data generating mechanism. When data are generated under the null hypothesis, P is the probability of type I error. When data are generated under an alternative hypothesis, *P* is the power of the test under that particular alternative. We estimate *P* by 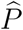, the proportion of the simulated datasets for which the test procedure rejects the null hypothesis (i.e., permutation test p-value < 0.05). For the test of localized differences we estimate the integrated actual coverage for pointwise confidence intervals (*IAC_p_*) and the integrated actual coverage for joint confidence intervals(*IAC_J_*) as described in Crainiceanu et al. (2012). Note that *IAC_P_* is based on the pointwise confidence intervals computed with the *z*-score cutoff. We also explore the pointwise and joint confidence intervals with cutoffs based on *t* distribution with (*n*/2 – 1) degrees of freedom and denote the corresponding integrated actual coverages as *IAC_Pt_* and *IAC_Jt_*, respectively. The estimates 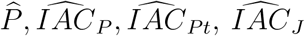 and 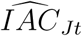 for the different scenarios are shown in Table 1.

**Table 1.**
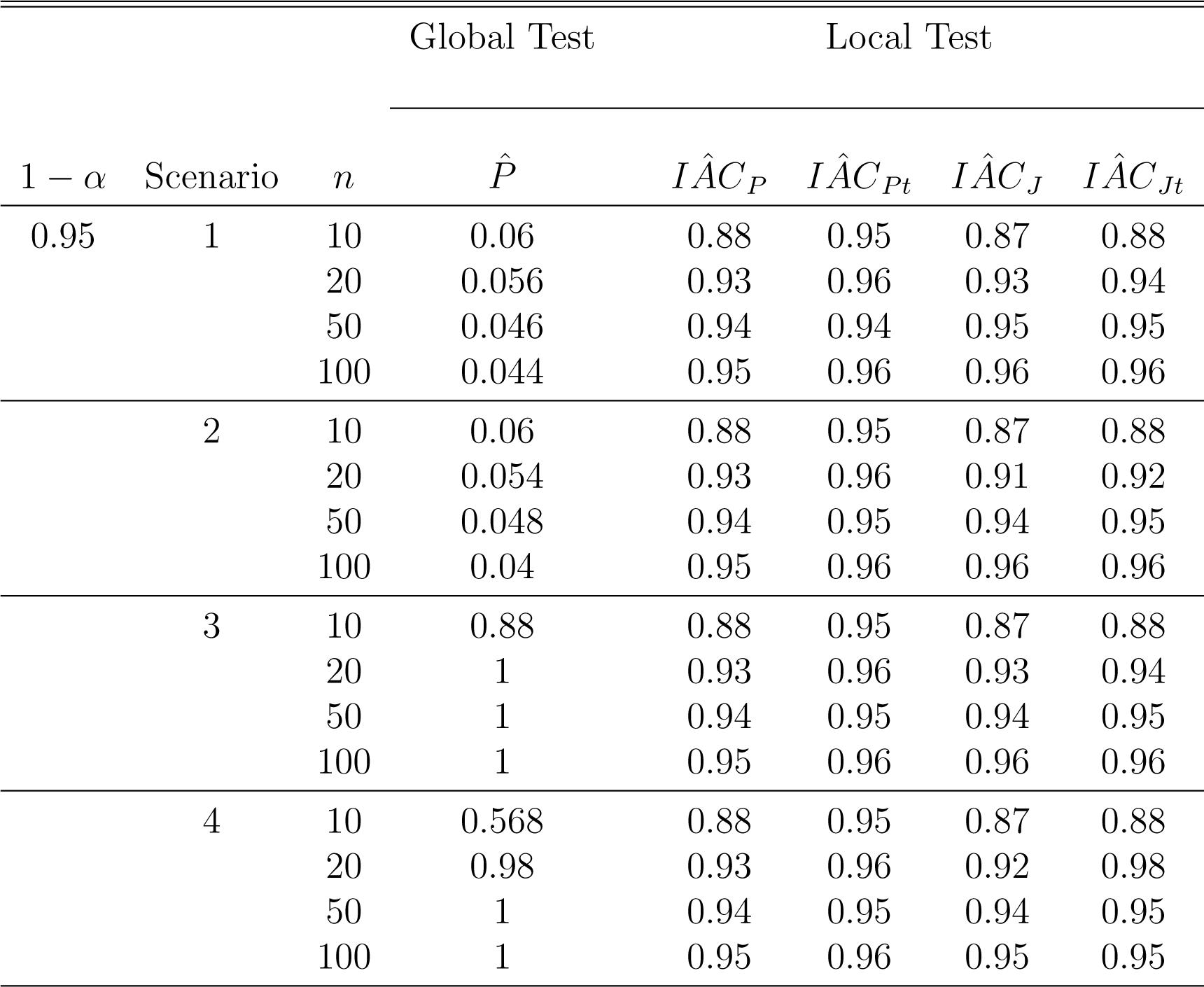
Simulation results: *Scenario 1 corresponds to β*_00_(*t*) = *β*_10_(*t*) = 0.91, *β*_01_(*t*) = *β*_11_(*t*) = (–0.77, 0.04); *Scenario 2 corresponds to β*_00_(*t*) = *β*_10_(*t*) = sin(*πt*),*β*_01_(*t*) = *β*_11_(*t*) = (–0.77, 0.04); *Scenario 3 corresponds to β*_00_(*t*) = 0.91, *β*_10_(*t*) = 1.40, *β*_01_(*t*) = *β*_11_(*t*) = (–0.77, 0.04); *Scenario 4 corresponds to β*_00_(*t*) = 0.1(1 +*t*)^2^, *β*_10_(*t*) = 0.5(1 – *t*)^2^, *β*_01_(*t*) = *β*_11_(*t*) = (–0.77, 0.04). *In all the scenarios, the two baseline covariates are generated independently in both the genotype groups from the same distribution, i.e., the empirical distribution of lean mass and fat mass to lean mass ratio in the control group.*

The global test produces the right Type I error for a sample size as little as *n* =10 as shown in the scenarios 1 and 2 in Table 1 where data are generated under *δ*(*t*) = 0. For the scenarios 3 and 4, where data are generated under *δ*(*t*) ≠ 0, the global test rejects the global null hypothesis in all the simulations for sample sizes greater than 20. For smaller sample sizes, e.g., n=10 and 20, the global test rejects the global null hypothesis in 57% and 98% of the 500 simulations, respectively.

For the 95% pointwise confidence intervals, if we use the cutoff based on *t* distribution with (*n*/2 — 1) degrees of freedom as opposed to the regular *z*-score cutoff, we get better coverages for the case *n* =10 under all scenarios. For higher sample sizes (i.e., *n* = 20, 50,100) the coverages with *t* distribution cutoff and *z*-score cutoff assume similar values and this finding is also consistent across all scenarios. The 95% joint confidence intervals suffer from undercoverage in the case *n* =10, but we see improvements in coverage for increased sample sizes.

## 4. Discussion

In this article, we provide simple and fast methods for testing if and where two covariate adjusted average curves are different.

One question we wanted to explore was whether there is an overall difference between the covariate adjusted curves. We develop a simple, easy-to-implement and novel test procedure by adapting the permutation test idea to functional outcomes. To the best of our knowledge, this is the first time such an approach is being proposed in the context of testing equality of two curves after covariate adjustment. We also propose a test for localized difference between the genotype groups (i.e., difference in average outcome at particular time points) using a non-parametric bootstrap of subjects (Crainiceanu et al., 2012). The methods in Crainiceanu et al. (2012) were developed for a matched case-control study. The major difference between our approach and the one presented in Crainiceanu et al. (2012) is that we are working with covariate adjusted curves.

The issue of covariate adjustment is of great importance in most scientific problems for reducing the variability and adjusting for baseline imbalances. Figures 2 and 4 show that the two genotype groups differ significantly with respect to the distribution of lean mass and fat mass to lean mass ratio. One of the strengths of our approach is that we perform the covariate adjustment in a statistically principled way: we first model the relationship of the oxygen consumption function and the two covariates: lean mass and fat mass to lean mass ratio using restricted maximum likelihood based functional regression methods; and then use the estimated regression functions and the within group sample averages of the covariates to estimate the within group average oxygen consumption. One issue of concern is that the empirical distribution of the fat mass to lean mass ratio has limited overlap between the genotype groups.

The global test shows good perfromance in simulation studies in terms of Type I error and power. The 95% pointwise confidene intervals, using z-score cutoff, suffer from under coverage for lower sample sized (e.g., n=10)., but the coverage improves with increasing sample size. Using cutoffs based on *t* distribution with (*n*/2 – 1) degrees of freedom leads to substantial improvement in coverage across all scenarios for *n* =10 and *n* = 20. The algorithm given in Crainiceanu et al. (2012) for computing 95% joint confidence intervals assumes multivariate normality of the difference function. We have explored other options, including assuming multivariate *t* distribution with (*n*/2 – 1) degrees of freedom, but they did not lead to significant improvement in coverage for lower sample sizes (e.g., n=10). In a future work, we plan to develop methods to address this issue.

We have also explored sensitivity of the developed methods to outliers. The exploratory analysis have identified one outlying animal in the IL-10^*tm*1*Cgn*^ group. We performed sensitivity analysis by excluding this animal. The results have some sensitivity to the exclusion of the outlying animal as indicated in the Appendix. Since the exclusion of the outlying animal impacts the findings both qualitatively and quantitatively, we decided to report both the results including and excluding the outlier. In such situations, it is recommended to include the outlying animal in the main analysis.

In summary, the key advantages of this method are its ease of implementation, efficiency and scalability. Although it is targeted to address the scientific question posed by the specific application, it can be adapted to a wide variety of biomedical and public healthsettings with similar design and data structure.

## Acknowledgements

This work was supported by the Older Americans Independence Center (OAIC) (Award Number: 2P30AG021334) funded by the NIH/National Institute of Aging. The statements and opinions in this article are solely the responsibility of the authors and do not necessarily represent the views of the Older Americans Independence Center or the NIH/National Institute of Aging. Conflict of Interest: None declared.

## Appendix

In this section, we show the additional results of sensitivity analysis after excluding the outlying animal from the dataset.

Figure 6 shows the point estimates and the 95% pointwise and joint confidence intervals for the difference in average oxygen consumption between the control mice and the IL-10^*tm*^ mice (excluding the outlying animal). When the oxygen consumption is normalized per gram of body weight (left panel) we observe a significant decrease in average oxygen consumption between early morning and late afternoon and again during late evening for the IL-10^*tm*^ mice compared to the control mice. The global geneotype effect is also significant (*p*-*value* = *0.006*). After adjustment for the lean mass and the fat mass to lean mass ratio, the localized genotype effect remains significant over certain smaller time intervals during afternoon and late night as indicated by the joint confidence intervals. As a consequence, the global genotype effect is also significant (*p*-*value* = *0.03*).

**Figure 6.**
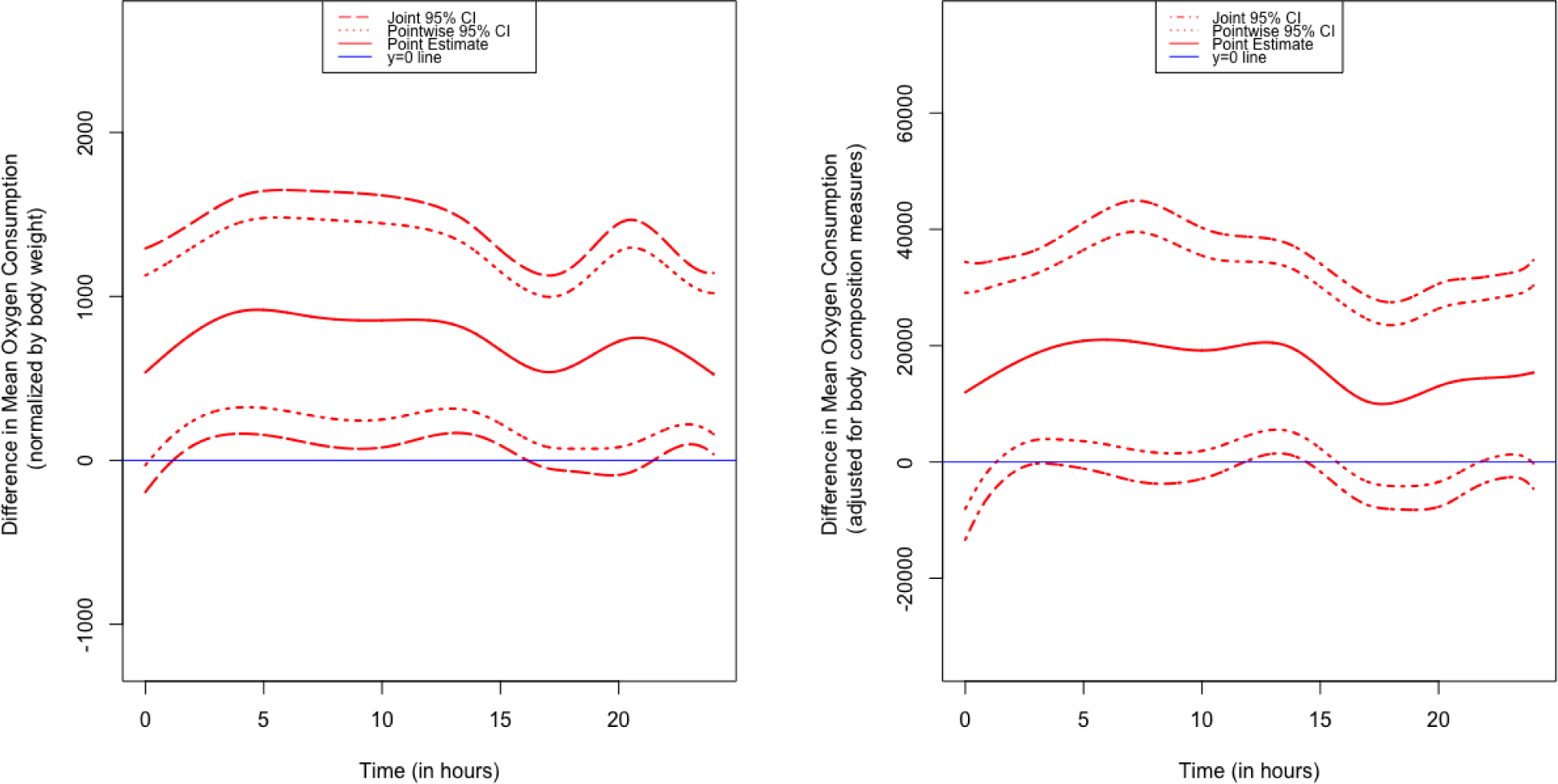
Results of sensitivity analysis after excluding the outlying animal in the IL-10^*tm*^ group. Plots showing 95% pointwise and joint confidence intervals for the difference in mean oxygen consumption between control mice and IL-10^*tm*^ mice: in the left panel the oxygen consumption is normalized by body weight (BW) and in the right panel the oxygen consumption is adjusted for the ratio of fat mass and lean mass (FM/LM) of the animal.

